# Mutations associated with pyrethroid resistance in Varroa mites, a parasite of honey bees, are widespread across the USA

**DOI:** 10.1101/2020.11.27.401927

**Authors:** Anabel Millán-Leiva, Óscar Marín, Krisztina Christmon, Dennis vanEngelsdorp, Joel González-Cabrera

## Abstract

**BACKGROUND:** Managed honey bees are key pollinators of many crops and play an essential role in the United States food production. For more than 10 years, beekeepers in the US have been reporting high rate of colony losses. One of the drivers of this colony loss is the parasitic mite *Varroa destructor*. Preserving healthy honey bee colonies in the US is dependent on a successful control of this mite. The pyrethroid *tau*-fluvalinate (Apistan^®^) was among the first synthetic varroacide registered in the US. With over 20 years of use, population of mites resistant to Apistan^®^ have emerged, and so it is unsurprising that treatment failures have been reported. Resistance in US mite populations to *tau*-fluvalinate is associated with point mutations at position 925 of the voltage-gated sodium channel, L925I and L925M.

**RESULTS:** Here, we have generated a distribution map of pyrethroid resistance alleles in Varroa samples collected from US apiaries in 2016 and 2017, using a high throughput allelic discrimination assay based on TaqMan^®^. Our results evidence that these *kdr*-type mutations are widely distributed in Varroa populations across the country showing high variability among apiaries.

**CONCLUSION:** We highlight the relevance of monitoring the resistance in mite populations to achieve an efficient control of this pest, and the benefit of implementing this methodology in pest management programs for varroosis.

## 1. INTRODUCTION

Beekeepers in the United States (US) have noted higher than acceptable annual colony losses in the past 14 years^1, 2^. Prior to 2006, colony losses in the US were estimated to average 5–10% each year^3^, but they significantly increased to average around 39% annually in the last 10 years BIP,^1^. These losses can be recovered by replacing the colonies, but it has an economic toll and extra management efforts that are estimated to be millions of US dollars per year^3–6^. These high levels of losses are a concern for US beekeepers, the farmers that relies on beekeepers to provide bees for pollination, and the consumers who benefit from the diversity of food products that honey bee pollination facilitates^6–9^.

Beekeepers and researchers seem to agree that there are different factors, some of them interacting synergistically, that affect the bee health^10–13^. Of these factors, there is mounting evidence that the largest single driver of colony losses is the parasite *Varroa destructor* and the viruses this mite vectors^10, 13–24^. The mite *V. destructor*^25^ is an obligated ectoparasite which feeds on the fat body of immature and adult honeybees^23^ weakening them by reducing their hosts immune response, changing their physiology, and shortening their lifespan^26–31^. Varroa show an extraordinary capacity for vectoring honey bee viruses, which in turn cause colonies to crash suddenly^11, 13^. Currently, Varroa causes more economic damage than all other bee diseases^9–11^.

Since untreated colonies experience a rapid reduction in health, regular treatment against Varroa mite have become an essential part of bee management for US beekeepers. The use of synthetic varroacides like the formamidine amitraz, the organophosphate coumaphos, and the pyrethroids flumethrin and *tau*-fluvalinate have long been in beekeepers “toolbox”^8, 32^. Although there are other non-synthetic acaricides and management techniques available, synthetic miticides are often the preferred choice because, when effective, they remove mites rapidly, do not cause obvious damage to bee populations, and are relatively cheap and easy to use^5, 32–34^.

Synthetic varroacides are not, however, always effective in reducing colony loss rates. Some mite populations have become resistant to specific compounds. As with other arthropod pests, intensive treatment schemes that rely exclusively on a single compound over consecutive seasons has resulted in the evolution of resistance in mite populations^35–39^. Currently, resistance to the three classes of synthetic compounds used for Varroa control has been reported^37, 40–42^.

In 1987, the pyrethroid *tau*-fluvalinate was approved for use in US beehives in response to the recent introduction of Varroa^43^. This product was a mainstay of mite management until the late 1990s, when mite resistance was widely observed^35–39^. In Varroa, the mechanisms of resistance to amitraz and coumaphos remain unclear, but resistance to pyrethroids has been correlated with three different point mutations in the voltage-gated sodium channel (VGSC), the main target for synthetic pyrethroids and DDT^44–46^. The substitution of certain amino acids in the VGSC sequence results in a reduced sensitivity to pyrethroids^47–49^. This is a common mechanism of resistance in arthropod species, usually known as *knockdown resistance* (*kdr*) or *super-kdr* depending on the specific amino acid substitution and the level of resistance recorded^44, 50–53^. In resistant Varroa populations from Europe, the mutation found associated with *kdr*-type resistance to pyrethroids is the substitution from the wild-type leucine to valine at position 925 of the channel protein (L925V) (numbered after the housefly *para-type* sodium channel protein). But in the US, *kdr*-type mutations in Varroa populations are associated with an isoleucine (L925I) or methionine (L925M) substitutions of the wild-type leucine at the same position^54–57^.

Early detection of mutations associated with resistance could be a useful tool for beekeepers wanting to prevent treatment failures. Surveys to monitor mutations associated with resistance have resulted in increased management success of pest species like mosquitoes, aphids, beetles and ticks^58–63^. These efforts take advantage of the several methodologies currently available for detecting resistant alleles in arthropod pests^47, 58, 64, 65^. In case of *V. destructor*, we have previously designed and tested a high throughput allelic discrimination assay based on TaqMan®. This method has been demonstrated to be robust, reliable and fast for genotyping large number of mites individually, hence able to determine the incidence of pyrethroid resistant and susceptible mites in a given population^54, 55, 66^.

Using this diagnostic tool on mites sampled in 2016 and 2017, we generated a distribution map of pyrethroid resistance mutations across the US. By examining changes in mutation prevalence over the two years in study, we were able to infer management practice recommendations that may be useful to beekeepers wishing to use an integrated pest management strategy to control Varroa populations.

## 2. MATERIALS AND METHODS

### 2.1 *Varroa destructor* samples

As part of the US National Honey Bee Disease Survey (NHBDS), female adult Varroa mites were collected from apiary level aggregate samples across the US in 2016 and 2017^67^. In brief, samples were collected by apiary inspectors or professional beekeepers in all participating states, included ¼ cup of bees scooped from each of 8 colonies in the same apiary. When possible, collected bees were scooped from a brood nest frame containing open and closed brood cells. The mites were dislodged from the bees using a soapy water shaker (adapted from Rinderer, De Guzman and Sylvester^68^). Collected and counted mites were placed into a 1.5 ml microcentrifuge tube containing 70% ethanol and shipped to the University of Valencia, Spain for allele frequency determination.

### 2.2 TaqMan^®^ diagnostic assays

A high throughput allelic discrimination assay based on TaqMan® was used to genotype individual mites from 118 and 109 different apiaries sampled across the US in 2016 and 2017, respectively. The assay accurately discriminates among wild-type mites and those carrying the mutations L925I and L925M in the VGSC^55^.

Genomic DNA was extracted from 15 or 16 individual adult female mites per apiary using a modified alkaline hydrolysis method described previously by González-Cabrera, Davies, Field, Kennedy and Williamson^54^ Briefly, the mites were placed individually in each well of 96-well flat-bottom microtiter plates containing 20 μL of 0.25 M of NaOH. The mites were then ground using a multiple homogenizer (BA/MH96, Burkard Scientific Ltd., Uxbridge, U.K.)^69^. Subsequently, 20 μL of Neutralization buffer (125 mM HCl, 0.5% Triton X-100 and 125 mM Tris/HCl pH 8.0) was added to each well, and the plate was spun at 4,000 rpm for 5 min. The supernatant containing the DNA was transferred to a new 96-well PCR microplate and stored at −20 °C until used.

Primers and probes for the TaqMan® assay were designed using Primer Express™ Software v.2.0 (Life Technologies) against the relevant *V. destructor* genomic sequence (Accession KC152655), as described in González-Cabrera, Rodríguez-Vargas, Davies, Field, Schmehl, Ellis, Krieger and Williamson^55^. Primers were designed to amplify a single 97 bp fragment flanking the position 925 of the VGSC (Forward primer: 5’-CCAAGTCATGGCCAACGTT-3’, Reverse primer AAGATGATAATTCCCAACACAAAGG-3’)^54^.

Three probes were used for the detection of the different alleles at position 925 of the VGSC. They were labelled with a different fluorophore at the 5’-end. Thus, the probe specific for the wild-type allele (5’-TTACCCAGAGCTCC-3’) was labelled with VIC®, the probe specific for the L925I mutation (5’-AGGTTACCTATAGCTCC-3’) was labelled with 6-FAM™, and the probe specific for the L925M (5’-TTACCCATAGCTCCTATC-3’) mutation was labelled with NED®. All probes have a non-fluorescent quencher (NFQ) and a minor groove binder (MGB) attached to the 3’-end to improve the allele discrimination accuracy^70^.

TaqMan assay mixture contained 1.5 μL of genomic DNA, 7.5 μL of 2x TaqMan® Fast Advanced Master Mix (Applied Biosystems), 0.9 μM of each primer and 0.2 μM of each probe, in a total volume of 15 μL. Reactions were carried out on a StepOnePlus™ Real-Time PCR system (Applied Biosystems), with the following cycling conditions: 2 min at 50 °C, 10 min at 95 °C, followed by 40 cycles of 15 s at 95 °C and 1 min at 63 °C, ending with a post-PCR read of 2 min at 50 °C. Increasing fluorescence in VIC® (538 nm excitation and 554 nm emission), 6-FAM^™^ (494 nm excitation and 518 emission) and NED^®^ (546 nm excitation and 575 emission) was monitored in real time. The results obtained were analyzed using StepOne Software v2.3 (Applied Biosystems).

Statistical differences in the distribution of genotypes and phenotypes were assessed by t-test using SPSS v25.0 (IBM SPSS Statistics, Armonk, NY: IBM Corp.) and the differences observed were considered significant if *P<0.05*. Multiple comparison ANOVA was used to analyze differences in the results grouped by state. For the comparison of data from the same apiary between the two years, statistical significance was calculated by Fisher’s exact test using GraphPad Prism 7 (GraphPad software, Inc, San Diego, CA).

## 3. RESULTS

In this study, a total of 3,576 mites collected from 228 apiaries across the US territory were genotyped for mutations at position 925 of the VGSC, the region associated with pyrethroid resistance in *V. destructor*^54, 55, 66^. We have quantified the allele variation for position 925 of this gene using a TaqMan^®^ assay and specific probes to detect the wild-type and resistant mutations reported in the US (L925I and L925M).

This method discriminates individual mite genotypes by comparing the intensity of fluorescence signals during each cycle of the PCR amplification process. Increment of fluorescence for only one dye indicates that the mite is homozygous for that allele, and intermediate increases in the fluorescence of two dyes indicates that the mite is heterozygous^55^. Samples in which no fluorescence increase was detected or with questionable results were Sanger sequenced to rule out the presence of an allele different from those analyzed. In this study, only L925 mutations previously found in the US were identified (L925I and L925M), so neither the L925V mutation nor any other mutation at position 925 were detected.

To allow comparisons, we normalized the data and present the results as a percentage of the population examined. General allele frequencies were calculated pooling data from apiaries across the country from each year of collection. The wild-type leucine was the predominant allele at position 925 of the VGSC for the global population of *V. destructor* in the US (54.7%), with no statistically significant differences between the two years (t test, *P>0.05*). For the two detected mutant alleles, the frequency of isoleucine was 25.1% in 2016 and 28.7% in 2017 and that of methionine 20.2% in 2016 and 16.6% in 2017. However, these differences were not statistically significant, between years or alleles (t test, *P>0.05*).

As it is possible to find the three alleles in the same colony (wild-type allele and the two resistant alleles), six different genotypes can be generated from their combination in female Varroa (L/L, I/I, M/M, L/I, L/M and I/M). *Kdr-type* resistance is inherited as a recessive trait^20, 45, 49^, then only mites with genotypes 925I/I, M/M and I/M will show the resistant phenotype. On the other hand, mites carrying at least one copy of the wild-type allele would be susceptible to pyrethroid treatment (genotypes 925L/L, L/I and L/M), as indicated by González-Cabrera, Rodríguez-Vargas, Davies, Field, Schmehl, Ellis, Krieger and Williamson^55^. The data reporting the genotypes and predicted phenotypes (Susceptible or Resistant to pyrethroids) for each sampled apiary are shown in the Supplementary Table 1.

Summarizing the results by year of collection, our data revealed that mites homozygous for the amino acid at position 925 were more frequent than heterozygotes, with a significant increase detected in homozygotes in 2017 (83.4 *vs* 87.9%, t-test, *P*=0.006). When considering all data, the susceptible homozygote (925L/L) was the most prevalent genotype (49.2 and 50.3% of total mites tested in 2016 and 2017, respectively), followed by the resistant homozygotes 925I/I (19.5 and 24.4% of total mites tested) and 925M/M (14.6 and 13.2%). The remaining three heterozygotes had frequencies below 5.8% (Fig. 1). A significant reduction in the overall proportion of mites 925L/M and I/M was observed between 2016 and 2017 (t test, *P*=0.039 and *P*=0.009, respectively); no other genotypes changed over these two years. Overall frequencies of susceptible and resistance phenotypes remained constant in these two years (Fig. 1).

**Figure 1.**
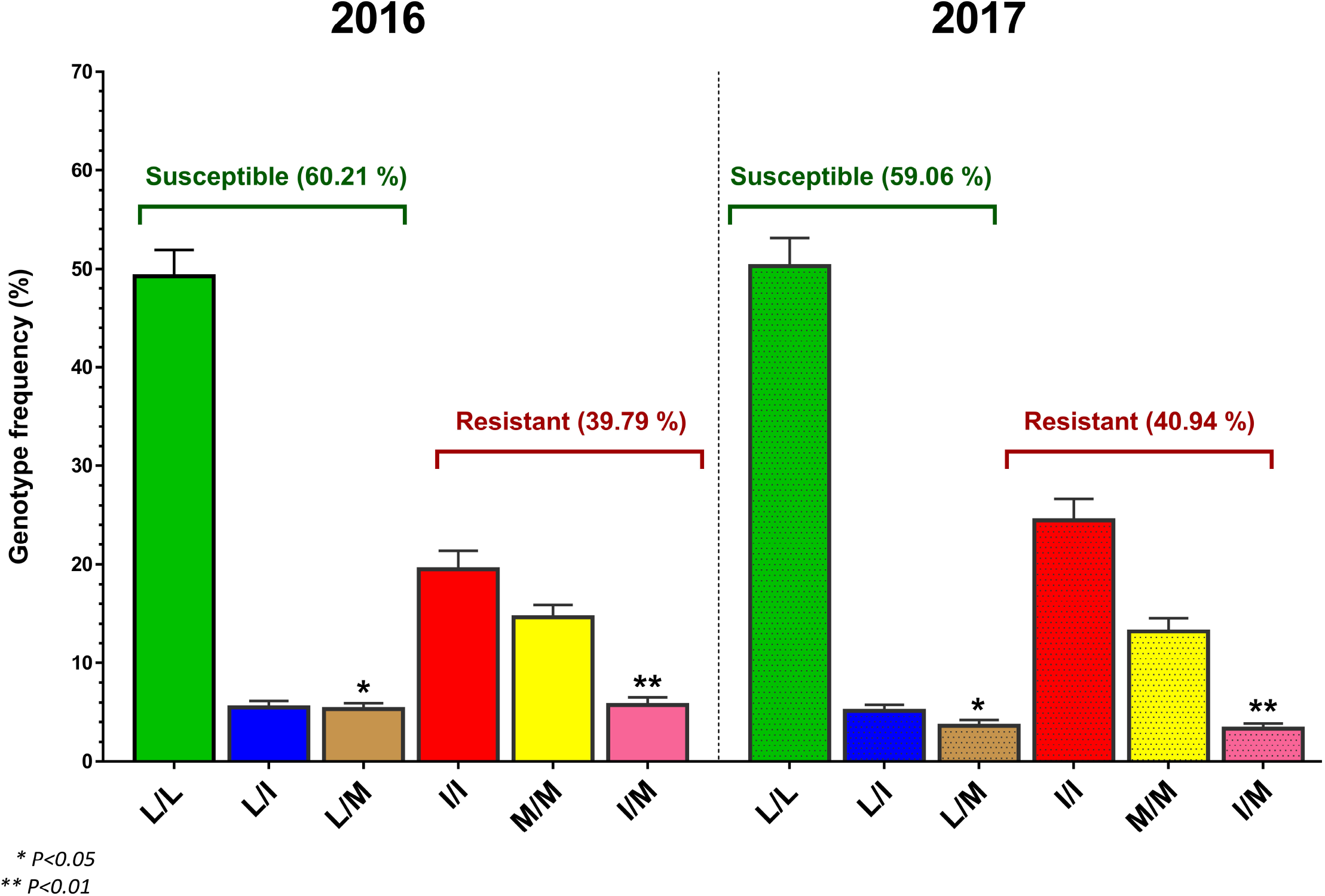
Genotype frequency of mutations at position 925 of the VGSC obtained from *V. destructor* collected across the US in 2016 and 2017, and predicted phenotype for pyrethroid resistance. L: wild-type L925, I: mutant L925I and M: mutant L925M.

The percentage of apiaries sampled in both years containing fully susceptible mites (i.e. genotypes L/L, L/I or L/M) was low and almost the same in both years, (mean ~9.5%). Worryingly, the proportion of apiaries completely ‘free’ of mites with resistant mutations was very low, (mean 2.2%).

To give spatial context to the analysis, overall genotype and phenotype frequencies were grouped by State for each year (Fig. 2, Supplementary Table 2). Our results showed that both mutant alleles are widespread throughout most apiaries in the country. The frequencies of genotypes and phenotypes among States, however, showed considerable variability. When comparing phenotype frequencies within individual states with the national average, we found that in 2016 seven out of 30 states differed (with higher rate of susceptible mites in Hawaii and Illinois; and lower in California, Georgia, Louisiana, North Dakota and Utah), and in 2017, nine out of 44 did (higher in Arizona, Puerto Rico and West Virginia, and lower in Colorado, New Hampshire, North Dakota, Oregon, and Wisconsin) (ANOVA, *P*<0.05) (Fig. 2, Supplementary Table 2).

**Figure 2.**
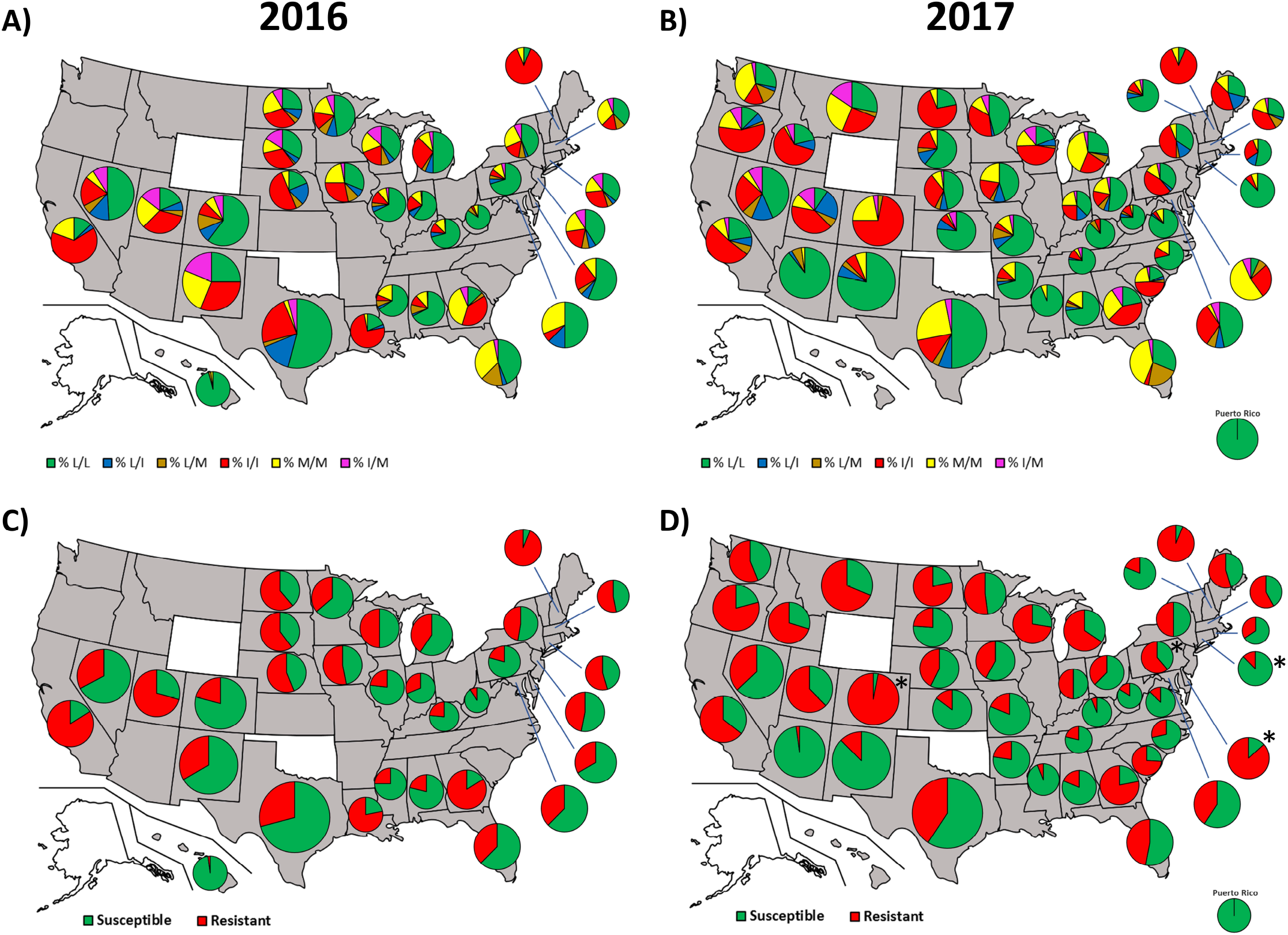
Distribution map of sampled *V. destructor*. Genotypes for the L925I and L925M mutations (A and B) and predicted Susceptible/Resistant to pyrethroids phenotype (C and D) by state for the years 2016 (A and C) and 2017 (B and D). Significant differences (t test, *P < 0.05*) between 2016 and 2017 are labeled (*).

Twenty five states were sampled in both years (Fig 3), of these only four (Colorado, Connecticut, Delaware and Pennsylvania) recorded changes in the frequency of susceptibility to pyrethroids (t test, *P*<0.05) (Fig. 2 and Fig. 3).

**Figure 3.**
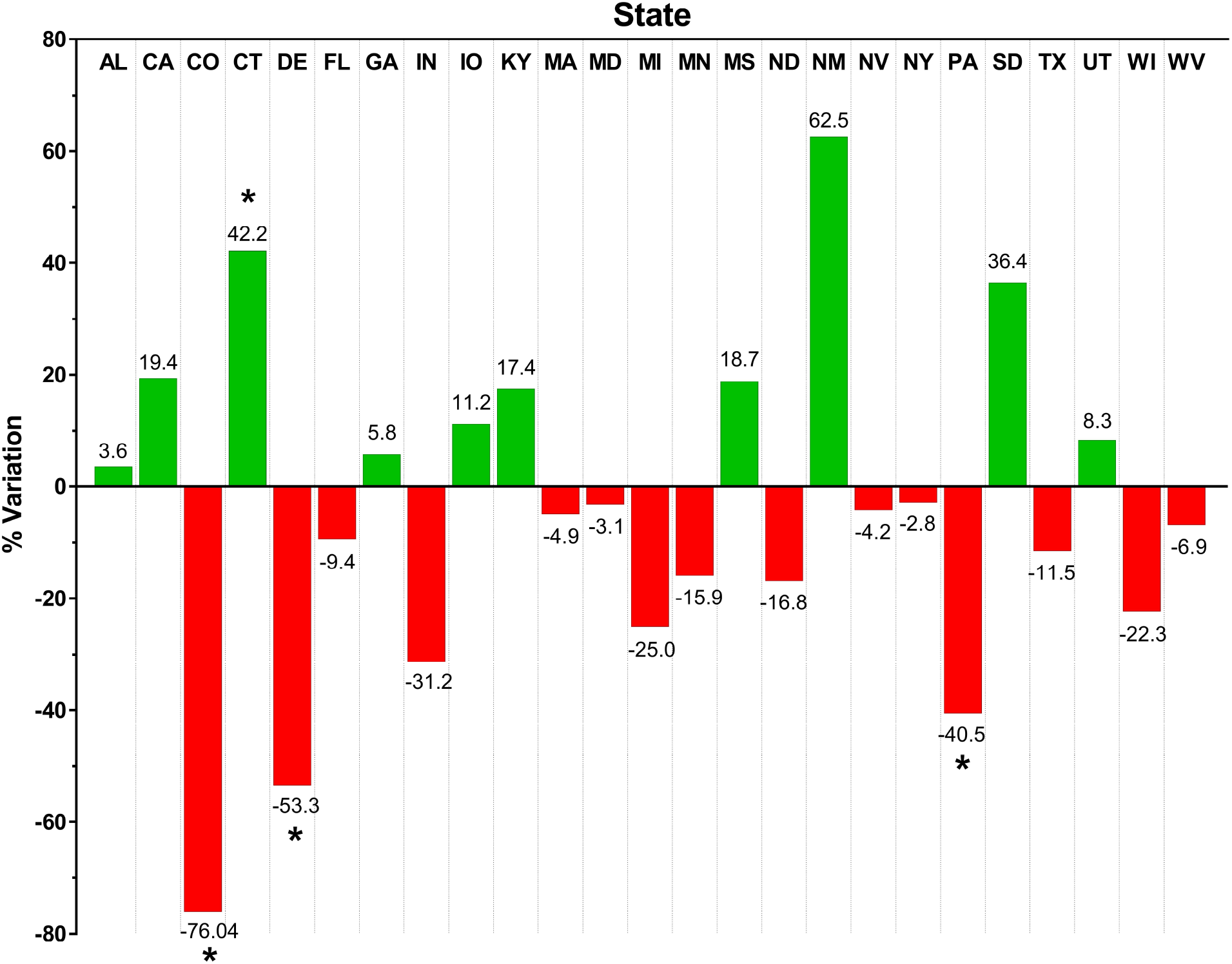
Variation between years 2016 and 2017 in the proportion of Varroa mites susceptible to pyrethroid treatment by State. Significant differences (t test, *P < 0.05*) between 2016 and 2017 are labeled (*).

Only eleven of the 110 samples from 2017 were collected in apiaries that were also sampled in 2016. The results of these apiaries in both years were compared side by side and are shown in Fig. 4 and Supplementary Table 3. This comparison showed that for seven of these apiaries the frequency of resistant mites differs between years (Fisher’s exact test, *P*<0.05), with six out of seven showing an increase in the proportion of resistant mites.

**Figure 4.**
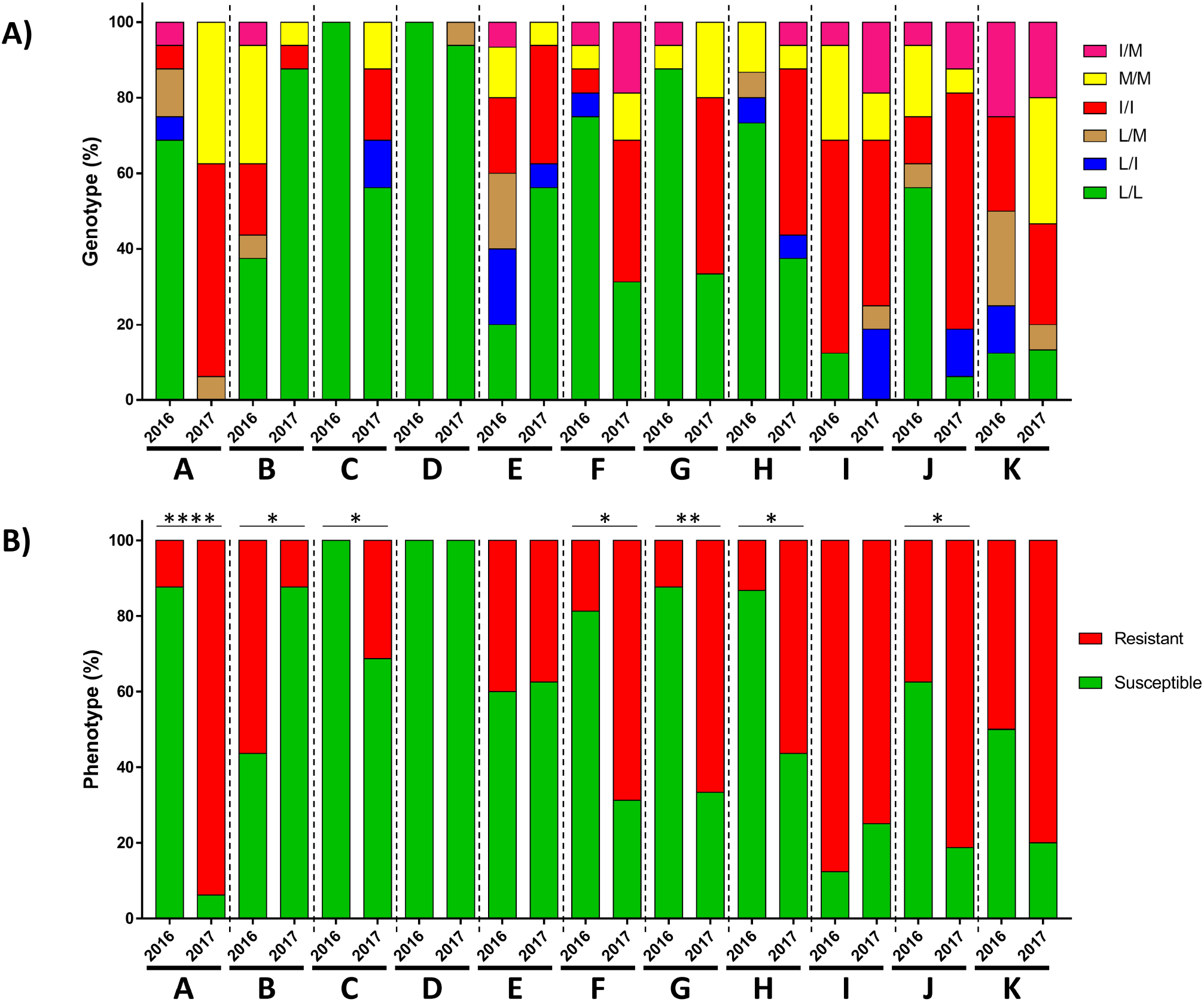
Genotype for position 925 of the *V. destructor* VGSC (A) and the associated pyrethroid susceptibility or resistance phenotype (B) in colonies from the same apiary sampled in 2016 and 2017. A to K letters refers to the apiary code. Significant differences between years are labelled as follows: Fisher’s exact test, \**P < 0.05, **P < 0.01, **** P < 0.0001*.

## 4. DISCUSSION AND CONCLUSIONS

In this study, we describe, for the first time, the frequency of pyrethroid resistant alleles in *V. destructor* populations across the US. The results over the two years surveyed suggest that the mutations responsible for pyrethroid resistance are widespread in the mite populations of US apiaries (Fig. 2).

Resistance to pyrethroids in *V. destructor* is associated with mutations at position 925 of the VGSC. The mutation L925V has been detected in European populations^54, 56, 57, 66, 71^ and in the US, the substitutions reported were L925I and L925M^55^. Our results confirm the presence of only these two mutations in the mites collected across the US. We also show that both alleles are widespread, with high incidence in some States (Fig. 2). This indicates that the ‘general’ use of *tau*-fluvalinate or other pyrethroid based products will have limited efficacy. These products, however, could have utility if they were included into an Integrated Pest Management strategy that monitors the frequency of resistant mites and promoted the appropriate Varroa control management approaches.

Varroa mite populations are highly inbreed, due to full sibling mating inside the brood cell (adelphogamy) along with haplo-diploid sex determination^32^. This biology makes heterozygotes very rare, and may favor the fixation of resistant alleles in the populations^72^. Our data supports this. Indeed, the prevalence of resistant genes in mites collected across the country were wide spread, which was facilitated, in all likelihood, by intensive use of the same treatment for many years^32^.

The overall proportion of resistant alleles and resistant phenotypes changed little between years (Fig. 1). However, a more in-depth analysis reveals that when the data is assessed by State and individual apiaries, significant changes over time were detected (Fig. 2, 3 and 4, Supplementary Table 2). This high variability suggests that allele frequencies are strongly influenced by beekeeper treatment strategies. Previously, the monitoring of the European resistant allele mutation, L925V, found a treatment effect on allele frequency^66^. Unfortunately, very few data were available regarding the treatment regime on the colonies sampled in both years. Access to the recent history of treatments would have been very valuable information to elucidate whether the changes in the frequencies were only influenced by the treatment regime or there are other factors contributing to the variation.

Some studies have evidenced the high incidence of mutations at position 925 in apiaries where pyrethroid treatment failed to control parasitism^54, 55, 66^. In contrast, in apiaries not treated with pyrethroids for some years, resistant mites are few or absent. In a recent study, González-Cabrera^66^ observed that for the European L925V mutation, the frequencies of resistant mites (homozygous 925V/V) decreased in a population when the treatment with pyrethroid was discontinued. Only one year of pyrethroid deprivation was enough to almost halve the resistant population. Similarly, Milani and Della Vedova^73^ reported a ten-fold decrease in resistant mites after 3 years without fluvalinate treatment. These studies combined, strongly suggest that the L925V mutation have a significant fitness penalty when the selective treatment is discontinued. Whether there is similar fitness penalty in mites with mutations L925I or L925M remains unknow, but it seems likely given that they are all at the same position of the channel protein. If so, it would bring some hope for developing IPM based strategies to control pyrethroid resistant Varroa in US apiaries.

A survey of beekeeping practices in US beekeeping operations, however, reported very low rate of *tau*-fluvalinate use among US beekeepers^5^, suggesting that there are other sources of selection pressure keeping resistant alleles at high frequencies in the US varroa population. One possible source is the accumulation of acaricide products in colony matrices like beebread and comb wax. *Tau*-fluvalinate is among the most prevalent and ubiquitous residue detected within colonies, largely because it is a lipophilic product and is sequestered in comb bees wax^74–76^. A recent survey of bee wax collected from US colonies reported that 70.8% of them had detectable levels of *tau*-fluvalinate (https://research.beeinformed.org/state_reports/pesticides_wax/). It is possible that exposure to accumulated lipophilic acaricides is exerting a continuous source of selection for resistant genes. Alternatively, selection pressure may be exerted through the contamination of bee forager collected food – nectar or pollen that was exposed to farmer applied products; in the US 37% of bee pollen samples collected contained fluvalinate, and 46% with residues of fluvalinate and other acaricides (Traynor K, Tosi S, Rennich K, Steinhauer N, Forsgren E, Rose R, Kunkel G, Madella S, Lopez D, Eversole H, Fahey R, Pettis J, Evans JD, vanEngelsdorp D, **Pesticides in Honey Bee Colonies: establishing a baseline for real world exposure over seven years in the USA-submitted for publication**). This suggests that appropriate comb management, may be a crucial component to long term varroa resistance management. Beekeepers may have some control over resistance in mite populations by ensuring their colonies are not chronically exposed to pyrethroid residues.

The data presented here is a picture of the situation in 2016 and 2017, but the level of resistance in a population may fluctuate over a short period of time depending on the treatment regime^66, 77^ or the exposure of mites to acaricide residues form previous treatments (see above). Since resistant alleles are ubiquitous, but not fixed, in the US mite population, proper actions could drive the frequency of resistance mutations to the minimum. By rotating varroacides with different modes of action, the use of fluvalinate in the US beekeeping operations may still have utility for longer-term strategies. Indeed, US commercial beekeepers who take a diverse approach to mite treatment lose fewer colonies (presumably because of increased mite treatment efficacy) than their less creative colleagues^5^. Predicting product efficacy based on allele frequencies data obtained using high-throughput DNA-based genotyping assays like that used here, would be a valuable tool for beekeepers making product use decisions.

One of the largest changes facing beekeepers today is controlling Varroa mites, a problem exacerbated by the limited number of control products available and the evolution of resistance in mite populations^8–32, 45, 78^. Although alternative non-synthetic based treatments are available, their efficacy is variable as outcomes are dependent on external factors, such as climatic and in-hive conditions and product application^13, 32, 79, 80^.

US beekeeers, specifically, have been suffering of high rate losses over the last 10 years. A national survey of honey bee diseases pointed the Varroa parasitism as the number one stressor for US operations and suggest that no less than 40% of US collonies had mite levels well above threshold, particularly in the fall when mite parasitism is most damaging^81^. Nonetheless, there is approximately 7% of US beekeepers that report not treating their hives for mites in any way^5^, which makes the incidence of mites and the risk of infection of nearby colonies higher. However, it is also clear that even for beekeepers using mite control methods, these are not always effective in reducing colony loss rates, partly because resistance to certain mite control products have evolved in the populations. In the near future the development of new effective varroacides is not very likely. Setting up precise and simple diagnostic tools for monitoring the presence of resistance in apiaries individually may become a critical approach for an effective and sustainable management of the mite in the short term. Furthermore, combination and rotation of varroacides based of different active ingredients would prevent from fixation of resistant alleles in mite population, ensuring the long-term efficacy for chemical and natural acaricides.

## 5. Acknowledgments

The authors want to thank all beekeepers and beekeeper associations for providing all mites samples used in this study.

## 6. Author contribution

AML and JGC conceived and designed research. DvE and KC contributed biological samples. AML and OM conducted experiments. AML analyzed data. AML and JGC wrote the manuscript. All authors revised and approved the manuscript.

## 7. Conflict of interest and declaration

The authors declare no competing interests

## Supplementary material

**Supplementary Table 1.** *Varroa destructor* 925 allele data obtained for each sampled apiary in absolute and normalized values.

**Supplementary Table 2.** Genotype and phenotype frequencies in the 925 mutation of the VGSC of *V. destructor* grouped by State for years 2016 and 2017.

**Supplementary Table 3.** Genotype and phenotype for the 925 mutation of the VGSC of *V. destructor* of mites from the eleven apiaries sampled in the consecutive years, 2016 and 2017 (Fisher’s exact, *P*<0.05).

## Notes

**Funding**, Joel González-Cabrera was supported by the Spanish Ministry of Economy and Competitiveness, Ramón y Cajal Program (grant: RYC-2013-261 13834). The work at the Universitat de València was funded by the Spanish Ministry of Economy and Competitiveness (grant: CGL2015-65025-R, MINECO/FEDER, UE), the Spanish Ministry of Science, Innovation and Universities (grant: RTI2018-095120-B-100). Sample collection in the USA was funded by the US National Honey Bee Disease Survey USDA-APHIS (16-8100-1624-CA, 15-8100-1624-CA).

### Competing Interest Statement

The authors have declared no competing interest.

