## Supplementary Table 2 for "Mutations associated with pyrethroid resistance in Varroa mites, a parasite of honey bees, are widespread across the USA"

**Supplementary Table 2.** Genotype and phenotype frequencies in the 925 mutation of the VGSC of *V. destructor* grouped by State for years 2016 and 2017.

|  |  | **Genotype** | | | | | | **Phenotype** | |
| --- | --- | --- | --- | --- | --- | --- | --- | --- | --- |
| **State** | **year** | **% L/L** | **% L/I** | **% L/M** | **% I/I** | **% M/M** | **% I/M** | **Susceptible** | **Resistant** |
| **Alabama** | **2016** | 65.18 | 0.89 | 11.61 | 12.50 | 8.93 | 0.89 | 77.68 | 22.32 |
|  | **2017** | 72.92 | 4.17 | 4.17 | 2.08 | 16.67 | 0.00 | 81.25 | 18.75 |
| **Arizona** | **2017** | 89.13 | 2.17 | 6.52 | 0.00 | 2.17 | 0.00 | 97.83 | 2.17 |
| **Arkansas** | **2017** | 70.97 | 3.23 | 3.23 | 9.68 | 12.90 | 0.00 | 77.42 | 22.58 |
| **California** | **2016** | 12.90 | 3.23 | 0.00 | 64.52 | 19.35 | 0.00 | 16.13 | 83.87 |
|  | **2017** | 22.58 | 6.45 | 6.45 | 51.61 | 9.68 | 3.23 | 35.48 | 64.52 |
| **Colorado** | **2016** | 60.42 | 8.33 | 10.42 | 8.33 | 6.25 | 6.25 | 79.17 | 20.83 |
|  | **2017** | 0.00 | 0.00 | 3.13 | 71.88 | 21.88 | 3.13 | 3.13 | 96.88 |
| **Connecticut** | **2016** | 36.00 | 4.00 | 5.33 | 28.00 | 16.00 | 10.67 | 45.33 | 54.67 |
|  | **2017** | 87.50 | 0.00 | 0.00 | 6.25 | 6.25 | 0.00 | 87.50 | 12.50 |
| **Delaware** | **2016** | 56.25 | 6.25 | 4.17 | 22.92 | 10.42 | 0.00 | 66.67 | 33.33 |
|  | **2017** | 6.67 | 0.00 | 6.67 | 26.67 | 53.33 | 6.67 | 13.33 | 86.67 |
| **Florida** | **2016** | 43.75 | 3.13 | 15.63 | 0.00 | 34.38 | 3.13 | 62.50 | 37.50 |
|  | **2017** | 31.25 | 0.00 | 21.88 | 3.13 | 40.63 | 3.13 | 53.13 | 46.88 |
| **Georgia** | **2016** | 12.90 | 0.00 | 3.23 | 38.71 | 38.71 | 6.45 | 16.13 | 83.87 |
|  | **2017** | 21.88 | 0.00 | 0.00 | 40.63 | 28.13 | 9.38 | 21.88 | 78.13 |
| **Hawaii** | **2016** | 95.74 | 0.00 | 2.13 | 0.00 | 2.13 | 0.00 | 97.87 | 2.13 |
| **Idaho** | **2017** | 21.28 | 8.51 | 0.00 | 61.70 | 2.13 | 6.38 | 29.79 | 70.21 |
| **Illinois** | **2016** | 67.74 | 5.16 | 3.87 | 12.26 | 8.39 | 2.58 | 76.77 | 23.23 |
| **Indiana** | **2016** | 75.00 | 4.17 | 2.08 | 12.50 | 6.25 | 0.00 | 81.25 | 18.75 |
|  | **2017** | 37.50 | 12.50 | 0.00 | 25.00 | 21.88 | 3.13 | 50.00 | 50.00 |
| **Iowa** | **2016** | 31.25 | 6.25 | 9.38 | 28.13 | 21.88 | 3.13 | 46.88 | 53.13 |
|  | **2017** | 45.16 | 9.68 | 3.23 | 19.35 | 22.58 | 0.00 | 58.06 | 41.94 |
| **Kansas** | **2017** | 76.60 | 8.51 | 0.00 | 6.38 | 2.13 | 6.38 | 85.11 | 14.89 |
| **Kentucky** | **2016** | 70.91 | 5.45 | 0.00 | 10.91 | 11.82 | 0.91 | 76.36 | 23.64 |
|  | **2017** | 87.50 | 0.00 | 6.25 | 6.25 | 0.00 | 0.00 | 93.75 | 6.25 |
| **Louisiana** | **2016** | 18.75 | 3.13 | 0.00 | 75.00 | 3.13 | 0.00 | 21.88 | 78.13 |
| **Maine** | **2017** | 29.03 | 16.13 | 0.00 | 41.94 | 12.90 | 0.00 | 45.16 | 54.84 |
| **Maryland** | **2016** | 50.00 | 12.50 | 0.00 | 6.25 | 31.25 | 0.00 | 62.50 | 37.50 |
|  | **2017** | 46.88 | 6.25 | 6.25 | 31.25 | 3.13 | 6.25 | 59.38 | 40.63 |
| **Massachusetts** | **2016** | 37.50 | 0.00 | 9.38 | 15.63 | 31.25 | 6.25 | 46.88 | 53.13 |
|  | **2017** | 29.03 | 3.23 | 9.68 | 38.71 | 16.13 | 3.23 | 41.94 | 58.06 |
| **Michigan** | **2016** | 50.00 | 6.25 | 3.13 | 28.13 | 9.38 | 3.13 | 59.38 | 40.63 |
|  | **2017** | 26.56 | 1.56 | 6.25 | 21.88 | 40.63 | 3.13 | 34.38 | 65.63 |
| **Minnesota** | **2016** | 47.87 | 8.51 | 7.45 | 13.83 | 15.96 | 6.38 | 63.83 | 36.17 |
|  | **2017** | 45.83 | 2.08 | 0.00 | 35.42 | 10.42 | 6.25 | 47.92 | 52.08 |
| **Mississippi** | **2016** | 65.63 | 3.13 | 6.25 | 6.25 | 18.75 | 0.00 | 75.00 | 25.00 |
|  | **2017** | 93.75 | 0.00 | 0.00 | 0.00 | 6.25 | 0.00 | 93.75 | 6.25 |
| **Missouri** | **2017** | 64.06 | 7.81 | 9.38 | 4.69 | 10.94 | 3.13 | 81.25 | 18.75 |
| **Montana** | **2017** | 28.13 | 0.00 | 3.13 | 25.00 | 28.13 | 15.63 | 31.25 | 68.75 |
| **Nebraska** | **2016** | 18.75 | 18.75 | 6.25 | 50.00 | 6.25 | 0.00 | 43.75 | 56.25 |
|  | **2017** | 46.81 | 6.38 | 4.26 | 31.91 | 8.51 | 2.13 | 57.45 | 42.55 |
| **Nevada** | **2016** | 49.09 | 12.73 | 5.45 | 18.18 | 5.45 | 9.09 | 67.27 | 32.73 |
|  | **2017** | 43.48 | 13.04 | 6.52 | 23.91 | 4.35 | 8.70 | 63.04 | 36.96 |
| **New Hampshire** | **2017** | 6.25 | 0.00 | 0.00 | 87.50 | 6.25 | 0.00 | 6.25 | 93.75 |
| **New Jersey** | **2016** | 41.05 | 5.79 | 6.32 | 19.47 | 18.42 | 8.95 | 53.16 | 46.84 |
| **New Mexico** | **2016** | 25.00 | 0.00 | 0.00 | 31.25 | 25.00 | 18.75 | 25.00 | 75.00 |
|  | **2017** | 78.13 | 6.25 | 3.13 | 6.25 | 6.25 | 0.00 | 87.50 | 12.50 |
| **New York** | **2016** | 43.52 | 2.78 | 6.48 | 15.74 | 24.07 | 7.41 | 52.78 | 47.22 |
|  | **2017** | 34.38 | 12.50 | 3.13 | 43.75 | 6.25 | 0.00 | 50.00 | 50.00 |
| **North Carolina** | **2017** | 71.43 | 0.00 | 0.00 | 12.70 | 15.87 | 0.00 | 71.43 | 28.57 |
| **North Dakota** | **2016** | 25.81 | 8.60 | 4.30 | 31.18 | 21.51 | 8.60 | 38.71 | 61.29 |
|  | **2017** | 21.88 | 0.00 | 0.00 | 71.88 | 6.25 | 0.00 | 21.88 | 78.13 |
| **Ohio** | **2017** | 51.56 | 4.69 | 6.25 | 15.63 | 18.75 | 3.13 | 62.50 | 37.50 |
| **Oregon** | **2017** | 12.50 | 6.25 | 2.08 | 56.25 | 14.58 | 8.33 | 20.83 | 79.17 |
| **Pennsylvania** | **2016** | 70.13 | 5.19 | 3.90 | 6.49 | 10.39 | 3.90 | 79.22 | 20.78 |
|  | **2017** | 35.48 | 3.23 | 0.00 | 45.16 | 12.90 | 3.23 | 38.71 | 61.29 |
| **Puerto Rico** | **2017** | 100.00 | 0.00 | 0.00 | 0.00 | 0.00 | 0.00 | 100.00 | 0.00 |
| **Rhode Island** | **2017** | 53.13 | 12.50 | 0.00 | 28.13 | 6.25 | 0.00 | 65.63 | 34.38 |
| **South Carolina** | **2017** | 19.35 | 3.23 | 3.23 | 48.39 | 22.58 | 3.23 | 25.81 | 74.19 |
| **South Dakota** | **2016** | 33.33 | 4.76 | 1.59 | 31.75 | 12.70 | 15.87 | 39.68 | 60.32 |
|  | **2017** | 60.87 | 10.87 | 4.35 | 17.39 | 6.52 | 0.00 | 76.09 | 23.91 |
| **Tennessee** | **2017** | 76.25 | 2.50 | 0.00 | 11.25 | 5.00 | 5.00 | 78.75 | 21.25 |
| **Texas** | **2016** | 54.17 | 14.58 | 2.08 | 22.92 | 2.08 | 4.17 | 70.83 | 29.17 |
|  | **2017** | 50.00 | 6.25 | 3.13 | 12.50 | 25.00 | 3.13 | 59.38 | 40.63 |
| **Utah** | **2016** | 18.75 | 6.25 | 4.17 | 33.33 | 22.92 | 14.58 | 29.17 | 70.83 |
|  | **2017** | 9.38 | 21.88 | 6.25 | 40.63 | 9.38 | 12.50 | 37.50 | 62.50 |
| **Vermont** | **2017** | 71.88 | 6.25 | 3.13 | 9.38 | 6.25 | 3.13 | 81.25 | 18.75 |
| **Virginia** | **2017** | 84.78 | 2.17 | 0.00 | 4.35 | 8.70 | 0.00 | 86.96 | 13.04 |
| **Washington** | **2017** | 28.13 | 3.13 | 12.50 | 15.63 | 37.50 | 3.13 | 43.75 | 56.25 |
| **West Virginia** | **2016** | 84.38 | 0.00 | 6.25 | 3.13 | 6.25 | 0.00 | 90.63 | 9.38 |
|  | **2017** | 75.00 | 5.00 | 3.75 | 12.50 | 3.75 | 0.00 | 83.75 | 16.25 |
| **Wisconsin** | **2016** | 38.10 | 4.76 | 7.14 | 19.05 | 17.86 | 13.10 | 50.00 | 50.00 |
|  | **2017** | 19.15 | 6.38 | 2.13 | 46.81 | 14.89 | 10.64 | 27.66 | 72.34 |
| **Overall US** | **2016** | 49.24 | 5.50 | 5.35 | 19.49 | 14.65 | 5.77 | 60.09 | 39.91 |
|  | **2017** | 50.26 | 5.17 | 3.63 | 24.44 | 13.17 | 3.33 | 59.06 | 40.94 |
