## Supplementary Table 3 for "Mutations associated with pyrethroid resistance in Varroa mites, a parasite of honey bees, are widespread across the USA"

**Supplementary Table 3.** Genotype and phenotype for the 925 mutation of the VGSC of *V. destructor* of mites from the eleven apiaries sampled in the consecutive years, 2016 and 2017 (Fisher’s exact, *P*<0.05).

|  |  | **Genotype** | | | | | | **Phenotype** | |
| --- | --- | --- | --- | --- | --- | --- | --- | --- | --- |
| **Beekeeper code** | **year** | **# L/L** | **# L/I** | **# L/M** | **# I/I** | **# M/M** | **# I/M** | **Susceptible** | **Resistant** |
| **CO-13** | **2016** | 11 | 1 | 2 | 1 | 0 | 1 | 14 | 2 |
|  | **2017** | 0 | 0 | 1 | 9 | 6 | 0 | 1 | 15 |
| **CT-06** | **2016** | 6 | 0 | 1 | 3 | 5 | 1 | 7 | 9 |
|  | **2017** | 14 | 0 | 0 | 1 | 1 | 0 | 14 | 2 |
| **IN-04** | **2016** | 16 | 0 | 0 | 0 | 0 | 0 | 16 | 0 |
|  | **2017** | 9 | 2 | 0 | 3 | 2 | 0 | 11 | 5 |
| **KY-06** | **2016** | 16 | 0 | 0 | 0 | 0 | 0 | 16 | 0 |
|  | **2017** | 15 | 0 | 1 | 0 | 0 | 0 | 16 | 0 |
| **MN-04** | **2016** | 3 | 3 | 3 | 3 | 2 | 1 | 9 | 6 |
|  | **2017** | 9 | 1 | 0 | 5 | 1 | 0 | 10 | 6 |
| **MN-12** | **2016** | 12 | 1 | 0 | 1 | 1 | 1 | 13 | 3 |
|  | **2017** | 5 | 0 | 0 | 6 | 2 | 3 | 5 | 11 |
| **PA-11** | **2016** | 14 | 0 | 0 | 0 | 1 | 1 | 14 | 2 |
|  | **2017** | 5 | 0 | 0 | 7 | 3 | 0 | 5 | 10 |
| **PA-12** | **2016** | 11 | 1 | 1 | 0 | 2 | 0 | 13 | 2 |
|  | **2017** | 6 | 1 | 0 | 7 | 1 | 1 | 7 | 9 |
| **UT-07** | **2016** | 2 | 0 | 0 | 9 | 4 | 1 | 2 | 14 |
|  | **2017** | 0 | 3 | 1 | 7 | 2 | 3 | 4 | 12 |
| **WI-17** | **2016** | 9 | 0 | 1 | 2 | 3 | 1 | 10 | 6 |
|  | **2017** | 1 | 2 | 0 | 10 | 1 | 2 | 3 | 13 |
| **WI-21** | **2016** | 1 | 1 | 2 | 2 | 0 | 2 | 4 | 4 |
|  | **2017** | 2 | 0 | 1 | 4 | 5 | 3 | 3 | 12 |
